# A pre-loaded and assembled human Ago2-miRISC orchestrates *Shigella*-induced vacuolar rupture through the targeting of RhoGDIα

**DOI:** 10.1101/2025.04.23.650266

**Authors:** L Schiavolin, D Filopon, M Bonnet, A Perez-Quintero, M Charvin, B Sperandio, PJ Sansonetti, G Tran Van Nhieu, L Navarro

## Abstract

The enteroinvasive bacterium *Shigella flexneri* causes bacillary dysentery. It invades intestinal epithelial cells and forms a bacterium-containing vacuole, which it rapidly ruptures to reach the cytosol. Here, we found that human Argonaute 2 (Ago2), the central component of miRNA-induced silencing complex (miRISC), positively regulates early steps of *Shigella* infection, including vacuolar rupture. This is consistent with the rapid recruitment of Ago2 at *Shigella* entry foci. The ability of Ago2 to form an assembled miRISC was found required for vacuolar maturation, while its slicer activity is dispensable for this process. Furthermore, we found that the RhoGTPase inhibitor RhoGDIα is targeted by specific miRNAs, pre-loaded in Ago2, which, collectively, accelerate vacuolar rupture. Therefore, a pre-loaded and assembled Ago2-miRISC orchestrates vacuolar maturation by targeting RhoGDIα. These findings may have wider implications in the understanding of how stored Ago2-loaded miRNAs control immediate steps of infection, prior to the differential expression of microbe-responsive miRNAs.

## INTRODUCTION

*Shigella flexneri* (*Shigella*) is the most common cause of bacillary dysentery worldwide, and is endemic in low-income countries^1^. It is also used as a model bacterium to dissect the mechanisms governing intracellular pathogenesis^2^. The life cycle of *Shigella* initially involves the capture of bacterial cells by filopodia, which are nanometre-thin membrane protrusions produced from the host cell^3,4^. *Shigella* further invades host cells through injection of effectors using a type III secretion system^3^. This process is associated with a massive cytoskeleton remodeling at the site of bacterial contact with the host cell membrane^5^. Such a reorganization of actin filaments forms membrane protrusions that are referred to as ‘actin foci’, which engulf *Shigella*^5,6^. *Shigella* triggers a local calcium response at these entry foci, which is detectable within minutes of bacterial contact with epithelial cells^7^. This effect stimulates a calpain-dependent proteolytic degradation of the E1 SEA2 enzyme^8^, which is presumably initiated at entry foci and further amplified through a sustained calcium response at a later phase of infection^7^. As a result, a global inhibition of sumoylation occurs in host cells, which increases the number of actin foci, further amplifying *Shigella* invasion^8^. These phenotypes are orchestrated by the depletion of a sumoylated version of RhoGDIα, which is tightly bound to Rho GTPases and extract them from the plasma membrane to keep them in an inactive state in the cytosol^9^. Therefore, the depletion of SUMO-RhoGDIα releases its negative effect over Rho GTPases, thereby promoting *Shigella* invasion. Once inside host cells, *Shigella* is embedded in a Bacterium-Containing Vacuole (BCV), which is rapidly ruptured and disintegrated for cytosolic access. Some molecular events controlling BCV maturation have been reported and include the formation and disassembly of an “actin cocoon” surrounding the BCV, and the recruitment of infection-associated macropinosomes (IAMs), which result from the resolution of membrane protrusions^10–13^. More specifically, *Shigella* recruits the exocyst machinery at IAMs through the GTPases Rab8/Rab11, a process leading to the clustering of IAMs around BCV^11^. The physical BCV-IAM membrane-membrane interaction further promotes IAM-mediated unwrapping of the ruptured vacuole membranes from *Shigella*, through a microtubule-dependent mechanism^14^. Such unpeeling of vacuolar membrane is also achieved through the *Shigella*-triggered subversion of Rab35, which is recruited in direct contact with the BCV^15^. Altogether, this series of molecular events is essential to ensure an effective *Shigella* vacuolar escape. *Shigella* further multiples in the cytosol. During this replication phase, it propels itself using actin-based motility^5^, a feature required to spread within cells but also in between cells. Importantly, a massive host transcriptional reprogramming is triggered during these steps of *Shigella* infection, which is orchestrated by intertwined host antibacterial defence and bacterial virulence responses^16,17^. Recent studies have unveiled a critical role of post-transcriptional gene regulatory processes in controlling host transcriptome reprogramming, notably through *Shigella*-responsive microRNAs (miRNAs)^4,18,19^.

MicroRNAs are central post-transcriptional regulators of gene expression. Upon Dicer-dependent processing of miRNA precursors, the resulting miRNA duplexes associate with Argonaute (Ago2) protein, the central component of the miRNA-induced silencing complex (miRISC). One selected strand, the guide, further triggers silencing of sequence complementary mRNA targets, through mRNA degradation and/or translational inhibition^20–23^. The phosphorylation of Ago2 at serine 387 triggers its binding to LIM domain-containing-protein 1 (LIMD1). LIMD1 then ensures a tight association between Ago2 and Trinucleotide repeat-containing adaptor 6A (TNRC6A), which binds to the PIWI domain of Ago2 through W-repeats (W-motifs)^24–27^. These molecular events are essential for High Molecular Weight (HMW)-miRISC assembly by recruiting downstream effector proteins, including those involved in decapping and deadenylation of mRNA^21,22,28–30^. This series of molecular events leads to the translational repression, destabilization and degradation of mRNA targets, and in turn regulates diverse biological processes. It is now well-established that a subset of individual human miRNAs accumulates differentially during bacterial infections and control host-bacteria interactions^31^. For example, the human microRNA miR-29b-2-5p promotes the formation of filopodia before cell contact, by targeting the UNC5 netrin-1 receptor *UNC5C*^4^. This phenomenon enhances the capture of *Shigella* by host cells and their subsequent invasion. In addition, this miRNA is degraded by the exonuclease PNPT1 at late timepoints of *Shigella* infection to modulate intracellular replication^4^. A miRNA triad has also been shown to dampen *Shigella* actin-based motility by repressing neuronal Wiskott-Aldrich syndrome protein (N-WASP), thereby altering its ability to spread intercellularly^19^. Two of these miRNAs, namely miR-4732-5p and miR-6073, were found induced during *Shigella* infection, likely as part of a host-triggered antibacterial defence response^19^. A recent report also provides evidence that miR-6762-5p is specifically expressed at late phase of *Shigella* infection and promotes RhoA-dependent cytoskeleton modifications to achieve successful intercellular spreading^18^. Although the role and regulation of individual human miRNAs have begun to be described in host-*Shigella* interaction, nothing is known about the dynamics and biological relevance of Ago2-miRISC during early steps of *Shigella* infection. Furthermore, the mechanisms by which miRNAs regulate their targets during immediate steps of bacterial infections remain elusive.

Here, we found that the Ago2-dependent miRNA pathway positively regulates *Shigella*-induced foci formation, invasion as well as vacuolar maturation. The latter function relies on the ability of Ago2 to form an assembled Ago2-miRISC, which is competent for miRNA-directed gene silencing. We did not find a differential regulation of miRNAs nor a differential accumulation of Ago2-bound miRNAs at an early infection timepoint, suggesting the involvement of a pre-loaded miRISC. Accordingly, a subset of miRNAs, pre-loaded in Ago2-miRISC, were found to orchestrate vacuolar rupture through the targeting of RhoGDIα. This study thus demonstrates a critical role of Ago2 in the dynamics of host-*Shigella* interactions and unveils a miRNA-dependent mechanism controlling early steps of *Shigella* infection.

## RESULTS

### Ago2 is recruited at *Shigella* entry sites and promotes foci formation, invasion and vacuolar rupture

To assess whether human Ago2 contributes to host-*Shigella* interaction, we first analyzed the dynamics of its subcellular localization during infection. More specifically, we transfected a functional GFP-Ago2 fusion in HeLa cells^32^, and further tracked GFP signal by live confocal microscopy after inoculation with the wild type *Shigella flexneri* M90T strain. Using this approach, we detected a rapid and transient recruitment of the GFP-Ago2 fusion at bacterial invasion sites (Figure 1A), which was concomitant with the onset of *Shigella* entry foci and followed the kinetics of actin foci formation^13,33^ (Figures S1A and S1B). By contrast, we did not observe such an enrichment of fluorescence signal at entry foci with an GFP-Rab4 fusion (Figures 1A and 1B), nor with the cytosolic vacuolar fluorescence marker Gal3-mOrange^34^ (Figure S1B), which served as negative controls. Based on these observations, we went on to investigate the possible biological function(s) of Ago2 during *Shigella* infection. For this purpose, we made use of a CRISPR/Cas9-based *Ago2*^-/-^ HeLa cell line^35,36^, which is deficient in Ago2 protein (Figure 1C). These cells were infected with *Shigella* and we further analysed the impact of a lack of Ago2 on different steps of infection. We found a reduced number of intracellular *Shigella* in *Ago2*^-/-^ *versus* control (CTL) cell lines (Figures 1D and S1C), suggesting that Ago2 promotes the capture of *Shigella* by host cells and/or its invasion within host cells. This observation is consistent with a role of miR-29b-2-5p in promoting *Shigella* capture by host cells and their subsequent invasion^4^. We further transfected the *Ago2*^-/-^ and CTL cells with GFP-Actin and Gal3-mOrange 24h prior *Shigella* infection, and monitored the kinetics of actin foci formation and cytosolic escape by measuring the time period between bacterial addition, foci onset and vacuolar rupture, respectively. A delay in foci formation was also observed in *Ago2*^-/-^ compared to CTL cells, which was mostly detected for foci occurring beyond ±17 min of *Shigella* addition to cells (Figures 1E and S1D). This suggests that Ago2 promotes foci formation, particularly during extended contact of bacteria with cells, which might contribute to its positive role on *Shigella* invasion (Figure 1D). We further found a substantial delay in vacuolar maturation in *Ago2*^- /-^ compared to CTL cells –median rupture time of 12.6 min *vs.* 8.3 min– which was more pronounced for late foci (Figures 1F and S1E), supporting a central role of Ago2 in vacuolar maturation, which we decided to investigate in more depth in the rest of this study.

**Figure 1.**
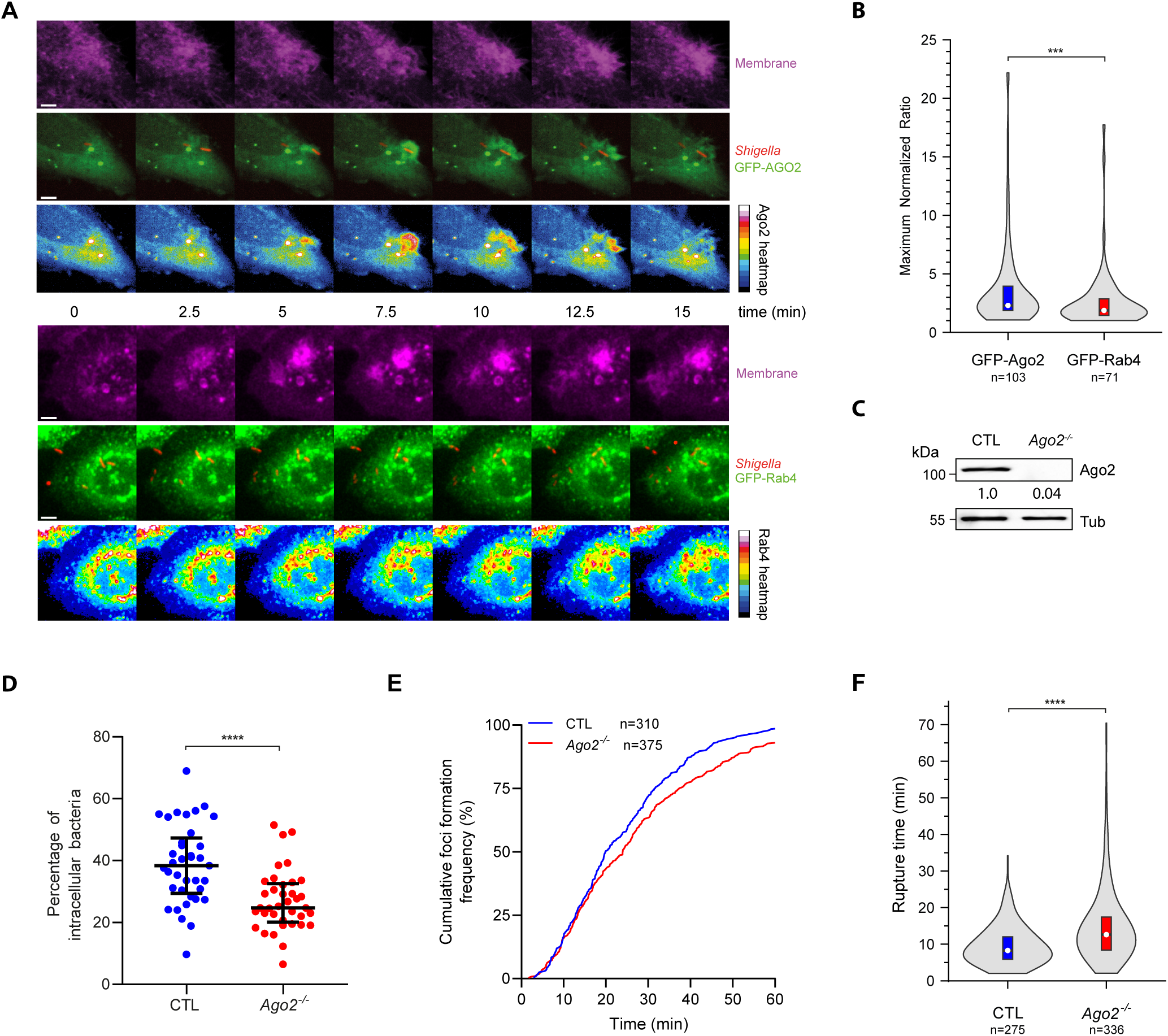
Ago2 is recruited at *Shigella* entry sites and promotes foci formation, invasion and vacuolar rupture. (A) Time-lapse fluorescent microscopy of *Shigella* mCherry (red) infecting either GFP-Ago2 (upper panel) or GFP-Rab4 (lower panel) expressing HeLa cells (green, middle). A deep red plasma membrane marker was used to monitor foci onset (purple, top). Heatmap of the pixel intensity of GFP-Ago2 at the indicated time points. The calibration bar for the heatmap is shown to the right (bottom). White arrows point to the focus formation (purple) and Ago2 signal at the focus (green). Scale bar, 5 µm. (B) Quantitative analysis of live-imaging data as illustrated in (A). The maximal recruitment of GFP-Ago2 or GFP-Rab4 (maximal fold increase of normalized fluorescence intensity) at *Shigella* entry foci was quantified. The data from at least three independent experiments are presented as violin plot. White circles show the medians, box limits indicate the 25^th^ and 75^th^ percentiles as determined by R software; violins represent density estimates of data and extend to extreme values. (C) Immunoblot analysis of HeLa control (CTL) and *Ago2^-/-^* cells. The α-Tubulin (Tub) was used as a loading control. Quantification of proteins accumulation is shown below the panel and is normalized with the loading control (Tub) and endogenous proteins level (CTL). (D) Quantification of intracellular *Shigella* expressed as a percentage of the total bacteria found attached to or inside HeLa control (CTL) and *Ago2^-/-^* cells. Medians, 25^th^ and 75^th^ percentiles are shown. Each dot represents data from a microscope field with at least 10 infected cells (n=37). Data is from at least three independent experiments. (E) Cumulative frequencies (up to 60 min) of foci formation time for *Shigella* infecting HeLa control (CTL) and *Ago2^-/-^* cells. (F) Vacuolar rupture time for *Shigella* in HeLa control (CTL) and *Ago2^-/-^* cells from panel (E) Same representation as in (B), with data from at least three independent experiments. Statistical tests: Mann-Whitney test (B, D, and F). *** p<0.001; **** p<0.0001.

### The Ago2-dependent miRNA pathway promotes the rapid rupture of *Shigella*-containing vacuoles

We next assessed whether the Ago2-dependent phenotype on vacuolar rupture could be dependent on the miRNA pathway. To test this, we monitored vacuolar rupture events in HeLa cells knocked-down for *Ago2*, the miRNA biogenesis factor *Drosha*^20,37^, the autophagy components *NDP52* and *ATG5*, which are required for the degradation of miRNA-free forms of human Ago2 and for miRNA functions^38^, or *P62*, a selective autophagy receptor that is not involved in Ago2 degradation nor in miRNA activity^38^, and thus served as a negative control. HeLa cells were silenced for 48h with siRNAs directed against each of these factors (Figures S2A and S2B), further transfected with Gal3-mOrange 24h prior infection. We then monitored the kinetics of vacuolar rupture. As expected, a delay in vacuolar rupture was detected in cells silenced for *Ago2* compared to cells transfected with control siRNAs –medians of 8.9 min and 6.6 min, respectively– (Figure 2A). When analysing the frequency distribution of vacuolar rupture events, we found that only ∼ 8 % of rupture events occurred within the first 4 min of infection in *Ago2* knocked-down cells compared to ∼ 23% in control cells (Figures 2B and 2C). This suggests that Ago2 starts exerting its positive effect on vacuolar maturation within the first minutes of infection, which is consistent with its rapid recruitment at entry foci (Figure 1A). As observed in *Ago2*-depleted cells, *Drosha* knocked-down cells displayed a retardation in the kinetics of vacuolar rupture events (Figures 2A and 2B). However, a milder effect over early rupture events was detected in *Drosha* knocked-down cells compared to *Ago2*-depleted cells, which was not found significantly different from control cells (Figure 2C). We also observed a delay in vacuolar rupture in the *NDP52* and *ATG5* knocked-down cells compared to control cells (Figures 2D and 2E), and the proportion of early rupture events was decreased and comparable to the one achieved in *Ago2* knocked-down cells (Figures 2E and 2F). By contrast, no phenotype was detected in cells depleted of *P62* (Figures 2D-2F). Altogether, these data demonstrate that the delayed vacuolar rupture events detected in *Ago2*-depleted cells are likely due to impaired miRNA functions manifested in these cells. Moreover, they provide evidence that the Ago2-dependent miRNA pathway contributes to the rapid rupture of *Shigella*-containing vacuoles.

**Figure 2.**
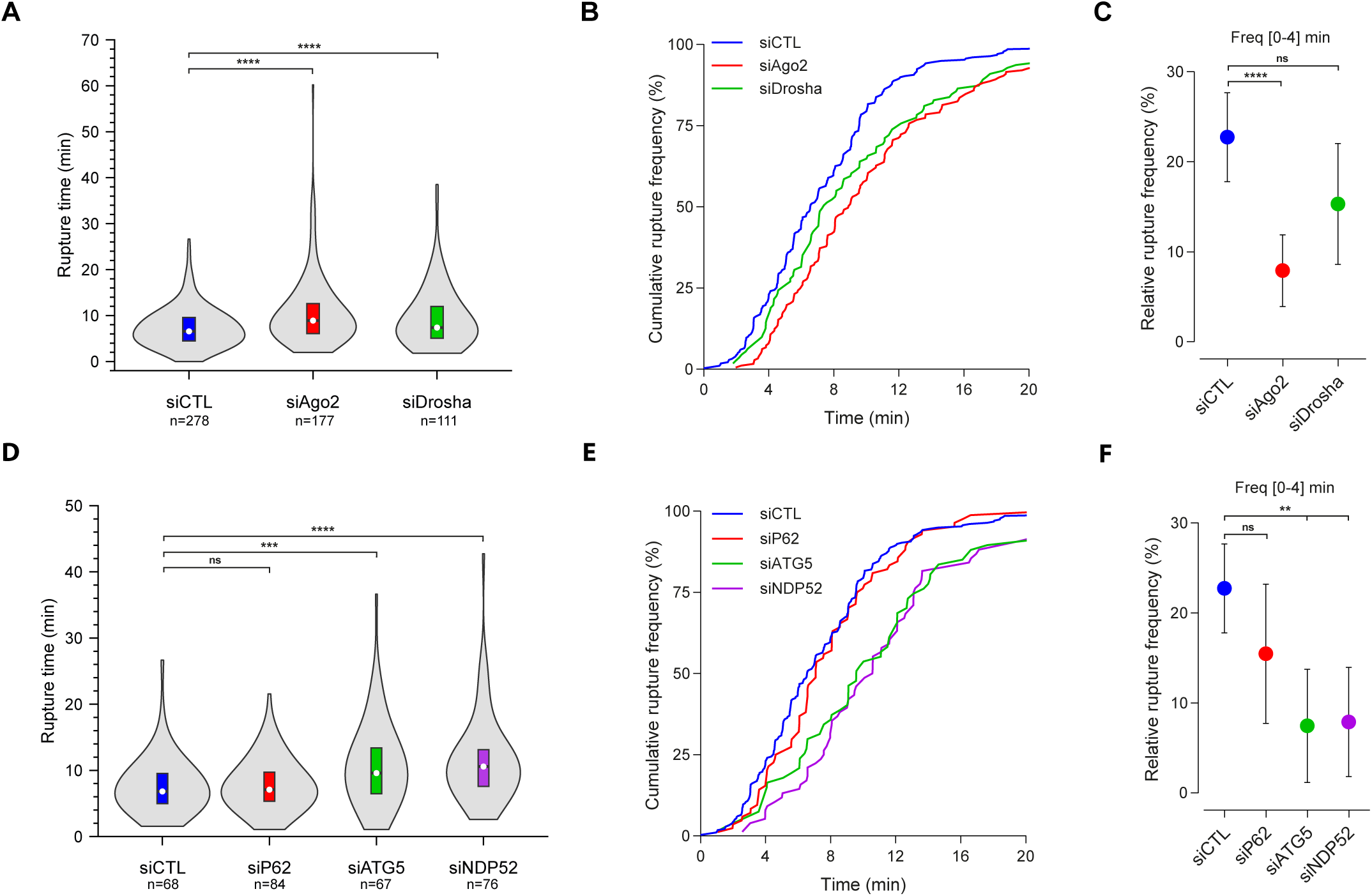
The Ago2-dependent miRNA pathway promotes the rapid rupture of *Shigella*-containing vacuoles. (A) Vacuole rupture time for *Shigella* in HeLa cells treated with control siRNAs (siCTL) or specific siRNAs against *Ago2* or *Drosha*. The data from at least three independent experiments are presented as violin plot. White circles show the medians, box limits indicate the 25^th^ and 75^th^ percentiles as determined by R software; violins represent density estimates of data and extend to extreme values. (B) Cumulative rupture frequencies (up to 20 min) of the data depicted in (A). (C) Relative frequencies with a 95% confidence interval (CI) of vacuolar rupture within the first four minutes after *Shigella* foci onset for the data presented in (A). (D) Vacuole rupture time for *Shigella* in HeLa cells knocked-down for *P62*, *ATG5* or *NDP52* using specific siRNAs. A non-targeting siRNA was used as a control condition (siCTL). Same representation as in (A), with data from at least three independent experiments. (E and F) Same representations as in (B) and (C) for data presented in (D). Statistical tests: Mann-Withney test (A and D); Chi-square (C and F). ns, non-significant; ** p<0.01; *** p<0.001; **** p<0.0001.

### The translational inhibition and mRNA degradation activities of Ago2 are critical for *Shigella*-induced vacuolar rupture, while its slicer activity is dispensable for this process

Ago2 possesses well-characterized silencing activities, including mRNA cleavage (“slicing”), translational inhibition and mRNA degradation. The two latter activities rely on the binding of TNRC6 proteins to specific residues of Ago2, which ensures Ago2-miRISC assembly^26,39^ (Figure 3A). To determine which of these Ago2 activities could be involved in *Shigella*-induced vacuolar rupture, we established a complementation assay in *Ago2*^-/-^ cells. Both the altered kinetics of vacuolar rupture and the frequency of rupture events manifested in these mutant cells were rescued when the GFP-Ago2 fusion was expressed *in trans* (Figures 3B-3D). The vacuolar rupture events occurred even faster than in CTL cells when a high concentration of GFP-Ago2 plasmid (*i.e.* 1 µg of plasmid, WT_1_ condition) was transfected in *Ago2*^-/-^ cells (Figures 3C and 3D), a phenomenon which was even detected within the first 4 min of infection (Figure 3E). Collectively, these data not only demonstrate that the GFP-Ago2 fusion effectively complement the delay in vacuolar rupture observed in *Ago2*^-/-^ cells, but also further support a positive role of Ago2 in promoting rapid vacuolar rupture. We further generated GFP-Ago2_D597A_ (slicer deficient^39^), GFP-Ago2_K660A_ (TNRC6A-binding deficient^40^) and GFP-Ago2_Y698A_ (TNRC6A binding-deficient^40^) constructs (Figure 3A), and tested them in the above complementation assay, upon transfection of 1 µg of each plasmid. Because the accumulation of the GFP-Ago2_Y698A_ mutant version was reduced compared to the other mutant versions (Figure 3B), we also transfected the GFP-Ago2 plasmid at a lower concentration (*i.e.* 0.5 µg of plasmid, WT_0.5_ condition). This allowed us to work in conditions expressing almost comparable protein levels of GFP-Ago2 and GFP-Ago2_Y698A_ (Figure 3B). The GFP-Ago2_D597A_ mutant version rescued a normal kinetics of vacuolar rupture events, to the same extent as the GFP-Ago2 construct (Figures 3C-3E). These data indicate that the slicer activity of Ago2 is dispensable for vacuolar rupture. By contrast, no complementation was observed upon expression of the stable GFP-Ago2_K660A_ and GFP-Ago2_Y698A_ mutants in *Ago2*^-/-^ cells (Figures 3C-3E). Altogether, these data indicate that the ability of Ago2 to interact with TNRC6 proteins, and thus to form an assembled Ago2-miRISC that is competent for miRNA-directed gene silencing, is essential for vacuolar rupture kinetics.

**Figure 3.**
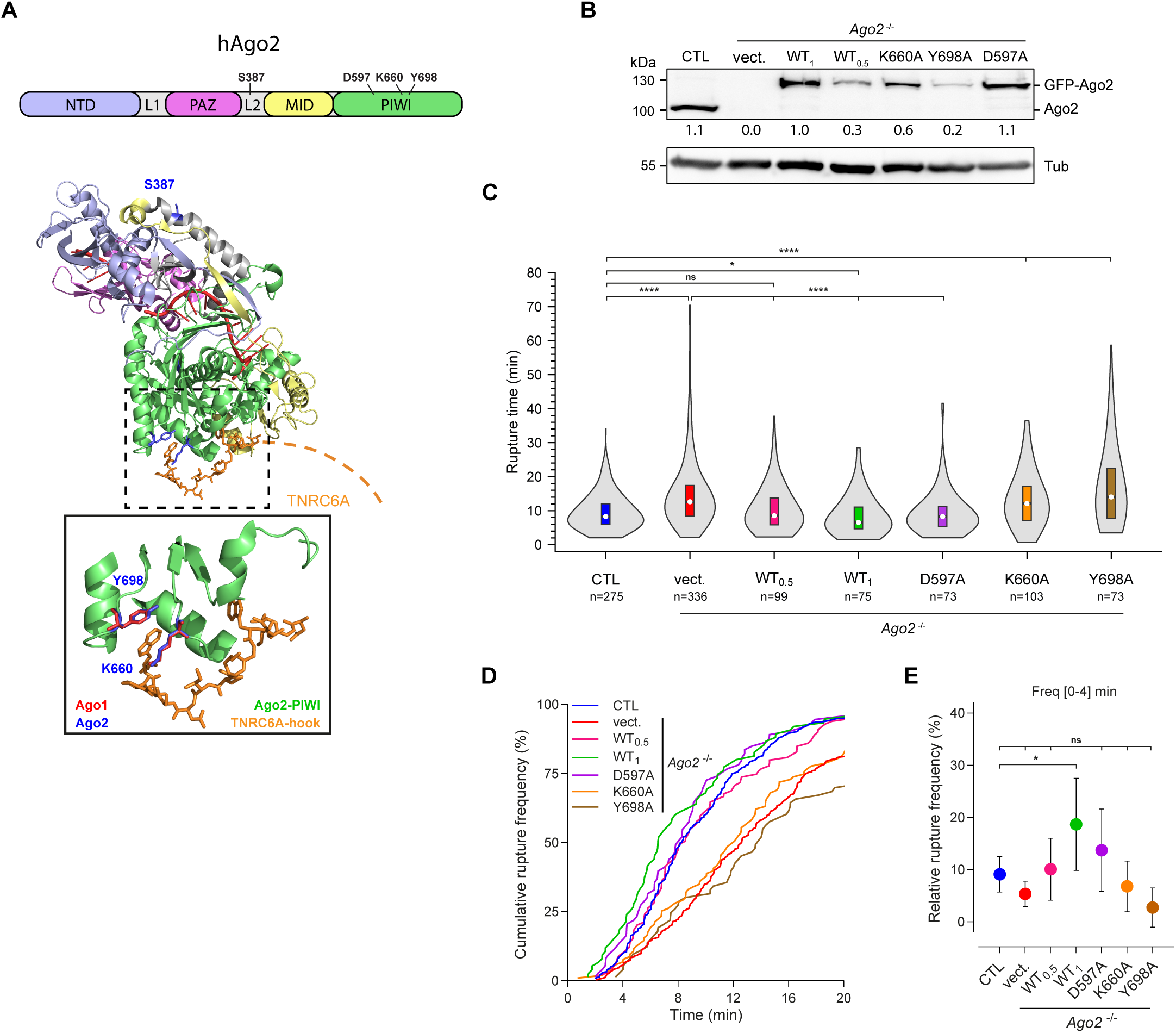
The translational inhibition and mRNA degradation activities of Ago2 are critical for *Shigella*-induced vacuolar rupture, while its slicer activity is dispensable for this process. (A) Domains organization of human Ago2 (hAgo2) and position of relevant residues (upper panel). The secondary structure of Ago2^23^ (PDB: 4F3T), coloured as in the upper panel, was superimposed to the structure of human Ago1-TNRC6A hook^40^ (PDB: 5W6V) using PyMol (RMS=1.73Å) (lower panel). The side chains of residues involved in Ago2/Ago1 interactions with TNRC6A (orange) are shown in the inset as blue and red, respectively. (B) Immunoblot analysis of HeLa control (CTL) and *Ago2^-/-^* cells transfected with 0.5 µg of GFP vector (vect.), 0.5 µg (WT_0.5_) and 1 µg (WT_1_) of GFP-Ago2 and 1 µg of GFP-Ago2 mutant variants as indicated. The α-Tubulin (Tub) was used as a loading control. Quantification of proteins accumulation is shown below the panel and is normalized with the loading control (Tub) and the level of GFP-Ago2 WT_1_. (C) Vacuole rupture time for *Shigella* in HeLa control (CTL), and *Ago2^-/-^* cells transfected with GFP (vect.) or GFP-Ago2 WT (at 0.5 or 1 µg) and mutant variants (1 µg). White circles show the medians, box limits indicate the 25^th^ and 75^th^ percentiles as determined by R software; violins represent density estimates of data and extend to extreme values from at least three independent experiments. (D) Cumulative rupture frequencies (up to 20 min) of data presented in (C). (E) Relative frequencies with 95% CI of vacuole rupture within the first four minutes after *Shigella* foci onset for data presented in (C). Statistical tests: Kruskal-Wallis (C), Chi-square (E). ns, non-significant; * p<0.05; **** p<0.0001.

### Both the phosphorylation of Ago2 at serine 387 and TNRC6 proteins, which orchestrate Ago2-miRISC assembly, positively regulate vacuolar rupture

The phosphorylation of Ago2 at serine 387 by Akt proteins entails Ago2-miRISC assembly, which is in turn competent for miRNA-directed translational inhibition and mRNA degradation^24,25^. By contrast, the unphosphorylated form of Ago2 at serine 387 is geared towards mRNA slicing activity^25,30^. To determine the relevance of this post-translational modification of Ago2 on vacuolar rupture, we generated and expressed a phospho-null GFP-Ago2_S387A_ or a phospho-mimic GFP-Ago2_S387E_ mutant constructs in *Ago2^-/-^* cells (Figure 4A). While the GFP-Ago2 and GFP-Ago2_S387E_ fully rescued the delayed rupture kinetics in *Ago2^-/-^* cells, the GFP-Ago2_S387A_ was not competent to do so (Figures 4B and 4C). Interestingly, we observed even faster and earlier rupture events upon expression of the GFP-Ago2_S387E_ compared to the GFP-Ago2, a phenomenon which was detectable within the first 4 min of infection (27.6% *vs.* 19.7%, p=0.1732) (Figure 4D). These data indicate that Ago2-S387 phosphorylation plays a key role in promoting the rapid rupture of *Shigella*-containing vacuoles. They also suggest that the downstream recruitment of TNRC6 proteins to Ago2, which is driven by this post-translational modification, might play a central role in vacuolar rupture. To test this, we further monitored vacuolar rupture in infected cells expressing an YFP-T6B fusion (Figures 4E-4G). T6B is a short TNRC6B-derived peptide, containing multiple Ago-interacting GW/WG dipeptides, which competes with TNRC6 proteins for Ago2 binding, thereby impairing miRNA functions^41^. A stable peptide mutated in all tryptophan residues (YFP-T6Bmut), which does not interact with Ago2^41^, and the YFP alone were used as negative controls. We found that expression of the YFP-T6B fusion triggered a significant delay in vacuolar rupture compared to YFP alone –median of 10 min *vs.* 6.5 min– (Figures 4F and 4G), supporting a role for TNRC6 proteins in vacuolar rupture. Conversely, infection of cells transfected with YFP-T6Bmut led to an overall rupture timing that was not different from cells transfected with YFP (Figures 4F and 4G). We also found that the depletion of TNRC6A from HeLa cells was sufficient to trigger a delay in the kinetics of vacuolar rupture events (Figures 4H-4J), although to a lesser extent than in the presence of T6B (Figures 4F and 4G). Altogether, these data provide evidence that both the Ago2-S387 phosphorylation and TNRC6 proteins, which orchestrate Ago2-miRISC assembly, are critical for rapid *Shigella*-induced vacuolar rupture.

**Figure 4.**
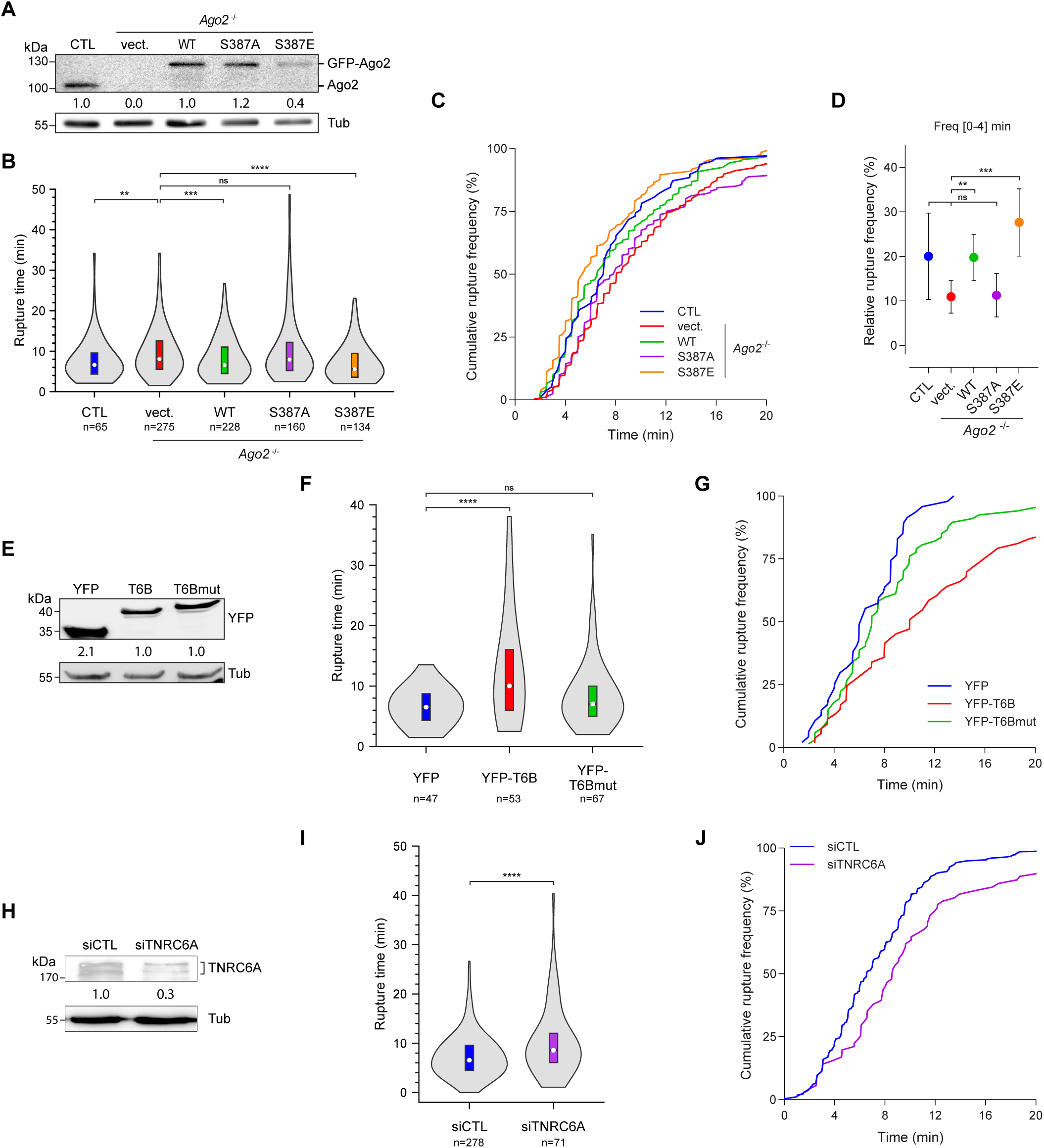
Both the phosphorylation of Ago2 at serine 387 and TNRC6 proteins, which orchestrate Ago2-mi-RISC assembly, positively regulate vacuolar rupture. (A) Level of endogenous Ago2 or transfected GFP-Ago2 variants in HeLa control (CTL) or *Ago2^-/-^* cells, respectively. (B) Vacuole rupture time for *Shigella* in HeLa control (CTL) or *Ago2^-/-^* cells transfected with GFP (vect.) or GFP-Ago2 WT and mutant variants as indicated. White circles show the median, box limits indicate the 25^th^ and 75^th^ percentiles as determined by R software; violins represent density estimates of data and extend to extreme values from at least three independent experiments. (C) Cumulative rupture frequencies (up to 20 min) of data depicted in (B). (D) Relative frequencies with 95% CI of vacuole rupture within the first four minutes after foci onset for data presented in (B). (E) Immunoblot analysis of HeLa cells transfected with YFP and YFP-T6B variants. (F) Vacuole rupture time for *Shigella* in HeLa cells expressing YFP, YFP-TNRC6B (T6B), or mutated YFP-TNRC6B (T6Bmut) peptides. Same representation as in (B), with data from at least three independent experiments. (G) Cumulative rupture frequencies (up to 20 min) of data in (F). (H) Immunoblot analysis of HeLa cells treated with the indicated siRNAs. (I) Vacuole rupture time for *Shigella* in HeLa cells treated with control siRNAs (siCTL) or siRNAs against *TNRC6A*. Same representation as in (B), with data from at least three independent experiments. (J) Cumulative rupture frequencies (up to 20min) of the data depicted in (I). Quantifications of proteins accumulation are shown below each panel and are normalized with the loading control (Tub) and endogenous or control proteins level (A, GFP-Ago2 WT; E, YFP-T6B; H, siCTL). Data presented in panel I and J were generated in the same experiments than data presented in Figure 2A. Statistical tests: Kruskal-Wallis (B), Mann-Whitney (F and I), Chi-square (D). ns, non-significant; ** p<0.01; *** p<0.001; **** p<0.0001.

### Thirteen Ago2-loaded miRNAs exhibit binding sites at the 3’ UTR region of *ARHGDIA*

To get some insights into the mechanism by which an assembled Ago2-miRISC directs *Shigella*-induced vacuolar rupture, we first retrieved the whole set of Ago2-loaded miRNAs at an early timepoint of infection. More specifically, we infected HeLa cells with *Shigella* for 15 min and further sequenced small RNAs from cell lysate (input) and Ago2-immunoprecipitated (RIP-Ago2) samples (Figure 5A). A Principal Component Analysis (PCA) showed clear separation between the Ago2-loaded fraction and the input, with no clear separation between mock and infected conditions within each fraction (Figure S3A). This analysis revealed an enrichment of 115 miRNAs in RIP-Ago2 fractions relative to inputs in the two independent experiments (Figure 5B and Table S1). We did not observe differences in the abundance of miRNAs between mock and infected conditions in both input and RIP-Ago2 samples (Figure 5B). These data imply that miRNAs are not differentially expressed, nor differentially loaded in Ago2, at this early timepoint of infection. They also suggest that a preloaded Ago2-miRISC is likely recruited at *Shigella* entry foci (Figure 1A), to promote rapid vacuolar rupture.

**Figure 5.**
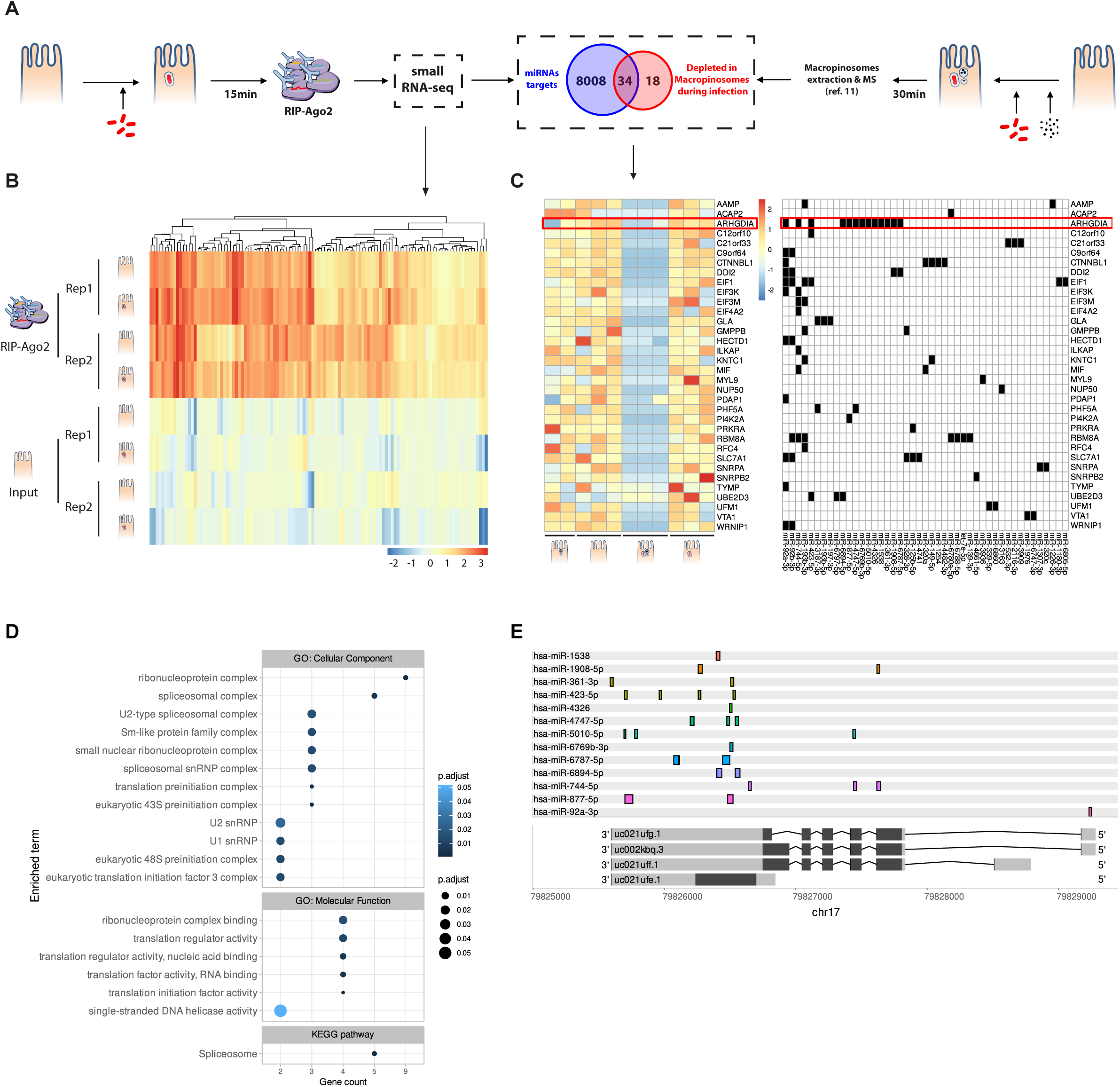
Thirteen Ago2-loaded miRNAs exhibit binding sites at the 3’ UTR region of *ARHGDIA*. (A) Schematic representations of both RNA immunoprecipitation (RIP) of Ago2 from *Shigella*-infected cells followed by sequencing of small RNAs (this study; left panel), and mass spectrometry (MS)-based analysis of the magnetically purified infection-associated macropinosomes (IAMs) of *Shigella*-infected cells^11^ (right panel). Integration of the two datasets is depicted in the central inset. (B) Heatmap of the 115 miRNAs enriched in RIP-Ago2 fraction compared to total small RNAs of uninfected and *Shigella*-infected cells (n=2). (C) Heatmap of the 34 candidate proteins associated to IAMs fraction compared to total proteins and depleted at IAMs during *Shigella* infection (left panel). Table depicting the absence (empty cell) or presence (filled cell) of binding site(s) for each of the 45 miRNAs targeting the mRNA of the 34 candidate proteins. *ARHGDIA* is highlighted by a red rectangle. (D) Enrichment analysis of KEGG pathways and gene ontology (GO) terms for the 8084 mRNAs that are targeted by the 115 Ago2-loaded miRNAs.

It has been reported that *Shigella* promotes the clustering of IAMs around these vacuoles, and that the physical interaction between these two compartments ensures a proper vacuolar maturation^10,11^. This phenomenon was shown to involve the recruitment of different factors at IAMs during *Shigella* infection^10,13^. Some of which were identified through a proteomic analysis from magnetically purified IAMs conducted at 30 min after *Shigella* infection^11^. By analysing this proteome dataset, we found that 14 peptides of Ago2 were specifically recovered in the magnetic *vs.* non-magnetic fractions of infected samples in the three independent experiments (Figure S3B). These data support an enrichment of Ago2 at IAMs at 30 min after *Shigella* infection. However, neither the number of IAMs per *Shigella* cell, nor their size distribution, were found altered in *Ago2*^-/-^ cells compared to control cells (Figures S3C and S3D). Therefore, Ago2 does not control the formation nor the size distribution of IAMs. It might rather exert its positive effect over vacuolar rupture by targeting negative regulators of this process at IAMs. To test this, we retrieved the whole set of experimentally validated targets of Ago2-loaded miRNAs^42^, and cross-referenced them with the list of proteins that were under-represented at IAMs (Figure 5A). We identified 34 candidates, which were found enriched in GO terms related to translational complexes and RNA-based regulatory complexes, and in KEGG pathways associated with spliceosome and RNA transport (Figures 5C and 5D). Among them, we found *ARHGDIA*, which encodes RhoGDIα, a negative regulator of RhoGTPases, which is known to promote *Shigella*-induced actin foci formation and invasion^8^. Significantly, we found that the *ARHGDIA* transcript was targeted by 13 different Ago2-loaded miRNAs, with a bias towards its 3’UTR region (Figures 5C and 5E). This observation is relevant because distinct miRNAs are known to act cooperatively to shut-down the expression of a given target more efficiently^43–45^. We also noticed that 6 out of the 13 Ago2-loaded miRNAs exhibit multiple binding sites at the *ARHGDIA* 3’UTR region (Figure 5E), which is another feature known to enhance target repression^43–45^. Based on these data, we decided to characterize the Ago2-miRISC-dependent regulation of *ARHGDIA* and its possible role in *Shigella*-induced vacuolar rupture.

### Ago2-miRISC positively regulates *Shigella*-induced vacuolar rupture by targeting RhoGDIα

We investigated whether RhoGDIα, and the targeting of its mRNAs by the above mentioned 13 miRNAs, could contribute to vacuolar rupture. To test this, we first expressed a CFP fusion of either the coding sequence, alone (GDIα) or with the miRNA-targeted 3’UTR region (GDIα + 3’UTR), of *ARHGDIA*, and further monitored vacuolar rupture. The expression of GDIα, which resulted in an enhanced RhoGDIα protein accumulation compared to the expression of the miRNA-targeted GDIα + 3’UTR version (Figure 6A), triggered a delay in vacuolar rupture compared to control cells expressing CFP alone (median of 9 min *vs.* 8 min) (Figures 6B and 6C). Furthermore, this phenomenon was already detected during the first 4 min of infection (Figure 6D), supporting a role for RhoGDIα in repressing rapid vacuolar rupture events. By contrast, expressing the miRNA-targeted GDIα + 3’UTR version in HeLa cells did not alter the kinetics of vacuolar rupture compared to control cells (Figures 6B-6D). Collectively, these data indicate that RhoGDIα acts as a negative regulator of vacuolar rupture. They also suggest that the targeting of the 3’UTR region of *ARHGDIA* mRNAs by miRNAs might limit the expression of RhoGDIα in HeLa cells, thereby reducing its negative effect over *Shigella*-induced vacuolar rupture. To characterize the Ago2-loaded miRNAs targeting the 3’UTR of *ARHGDIA* mRNAs, we further generated a miRNA sponge composed of a CFP construct carrying 3 binding sites for each of the 13 miRNAs targeting the *ARHGDIA* 3’UTR region (miRGDIα sponge, Figure 6E). As a negative control, we generated a CFP construct carrying unrelated miRNA binding sites (7xmiR21)^46^, referred to here as miRCTL sponge. We found a significant delay in vacuolar rupture in the presence of the miRGDIα sponge compared to the control miRCTL sponge –median of 10 min *vs.* 7.8 min for control cells– (Figures 6F and 6G). Furthermore, only 5.6% of the rupture events in the miRGDIα cells had occurred during the first 4 min compared to 15.5% in control cells expressing the miRCTL sponge (Figure 6H). Therefore, the miRNAs targeting the 3’UTR region of *ARHGDIA* mRNAs positively regulate vacuolar rupture and start exerting their functions within the first minutes of infection, which is consistent with a role of the Ago2-dependent miRNA pathway in this process (Figures 2A-2C and 3C-3E). Finally, we investigated whether depleting RhoGDIα could rescue the delay in vacuolar rupture observed in cells defective in miRNA functions. To this end, we knocked-down *ARHGDIA* in HeLa cells deprived of *Drosha* (Figure 6I), and further monitored the kinetics of vacuolar rupture. It is noteworthy that we did not perform this assay in the *Ago2*^-/-^ cells because exogenous siRNAs would unlikely be active in this background, given that Ago2 is the main silencing factor that directs siRNA-directed slicing of target transcripts in humans^39,47^. The delay in vacuolar rupture observed in the *Drosha*-depleted cells (median of 9 min) was fully rescued upon silencing of RhoGDIα in these miRNA-depleted cells (median of 5.5 min, Figures 6J and 6K). Furthermore, 32.6 % of the rupture events had occurred during the first 4 min in those conditions (Figure 6L), which is comparable to the effect detected upon expression of the GFP-Ago2_S387E_ version in HeLa cells (27.6 %, Figure 4D). These data demonstrate that RhoGDIα is causal for the delayed vacuolar rupture events observed in HeLa cells defective in miRNA functions. Altogether, these findings indicate that an assembled Ago2-miRISC, preloaded with 13 miRNAs, promotes vacuolar maturation by targeting RhoGDIα.

**Figure 6.**
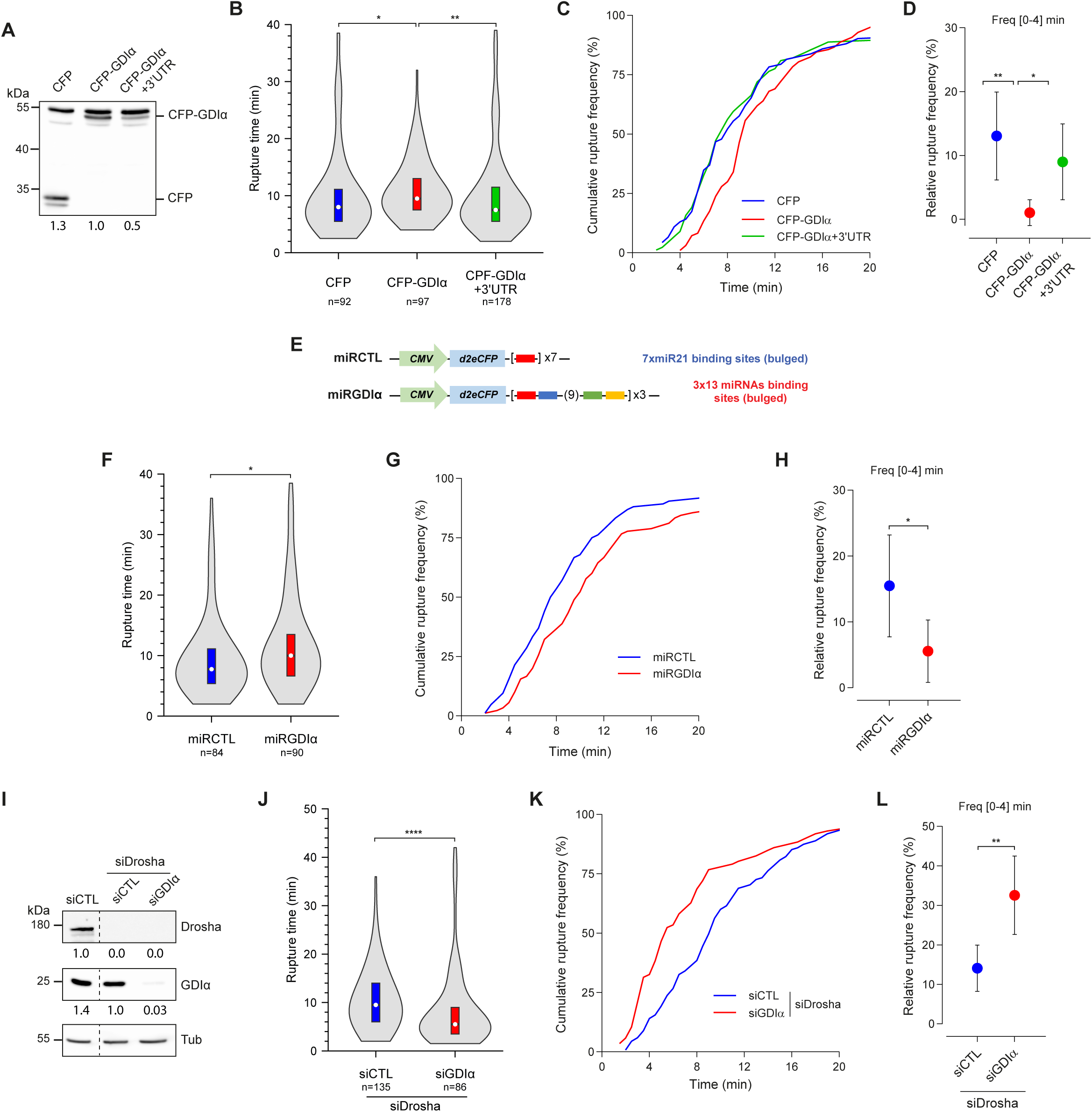
Ago2-miRISC positively regulates *Shigella*-induced vacuolar rupture by targeting RhoGDIα. (A) Immunoblot analysis of HeLa cells transfected with CFP or CFP-ARHGDIA with (GDIα + 3’UTR) or without (GDIα) its 3’UTR. The non-specific band around 55-kDa was used as a loading control. (B) Vacuole rupture time for *Shigella* in HeLa cells transfected with *ARHGDIA* expressing plasmids. White circles show the median, box limits indicate the 25^th^ and 75^th^ percentiles as determined by R software; violins represent density estimates of data and extend to extreme values from at least three independent experiments. (C) Cumulative rupture frequencies (up to 20min) of the data depicted in (B). (D) Relative frequencies with 95% CI of vacuole rupture within the first four minutes after *Shigella* foci onset for data presented in (B). (E) Detail of the sponge constructs for miRNAs. (F) Vacuole rupture time for *Shigella* in HeLa cells transfected with the CTL sponge (miRCTL) or the *ARHGDIA* sponge (miRGDIα). Same representation as in (B), with data from at least three independent experiments. (G and H) Same representations as in (C and D), respectively, for data presented in (F). (I) Immunoblot analysis of HeLa cells treated with the indicated siRNAs. The α-Tubulin (Tub) was used as a loading control. (J) Vacuole rupture time for *Shigella* in HeLa cells treated with both siRNAs against *Drosha* (siDrosha) and control siRNAs (siCTL) or specific siRNAs against *ARHGDIA* (siGDIα). Same representation as in (B), with data from at least three independent experiments. (K and L) Sames representations as in (C and D), respectively, for data presented in (J). Quantifications of proteins accumulation is shown below each panel and is normalized with the loading control (A, upper non-specific band; I, Tub) and endogenous or control proteins level (A, CFP-GDIα; I, siCTL). Statistical tests: Mann-Whitney (B, F, and J), Chi-square (D, H, and L). * p<0.05, ** p<0.01, **** p<0.0001.

## Discussion

Here, we showed that human Ago2 is recruited at *Shigella* entry foci within minutes of infection. This phenotype was found associated with a positive role of the Ago2-dependent miRNA pathway on early steps of infection, including vacuolar rupture. We identified 13 Ago2-loaded miRNAs targeting *ARHGDIA*, a novel miRNA target, whose cognate RhoGDIα protein levels was reduced at IAMs, and which negatively regulated vacuolar rupture (Figure 7). We also demonstrated that the delay in vacuolar rupture manifested in infected miRNA-defective cells is functionally linked to RhoGDIα. Altogether, these data suggest a model whereby *Shigella* subverts Ago2 activity at IAMs to silence RhoGDIα, thereby releasing its inhibitory effect towards RhoGTPases, such as Cdc42, which is known to orchestrate actin cocoon assembly for vacuolar maturation^12^. In this scenario, Ago2 would indirectly regulate *Shigella*-induced vacuolar rupture by promoting actin cage formation. Intriguingly, *Shigella* has previously been shown to deplete host cells from a sumoylated version of RhoGDIα on lysine 138 (K138), which presumably occurs at entry foci through the calcium/calpain-induced cleavage of the SUMO1 E1 enzyme SAE2^8^. Such sumoylation event on RhoGDIα enhances its binding activity to Rho GTPases, thereby restraining their biological activity^9^. As a result, a *Shigella*-induced clearance of SUMO-RhoGDIα favors the recruitment RhoGTPases at plasma membrane, which can in turn orchestrate actin foci formation and *Shigella* invasion^48,49^. Given that Ago2 was shown here to promote both steps of infection, we hypothesize that *Shigella* might additionally contribute to a miRNA-dependent depletion of RhoGDIα at entry foci. Such a dual regulatory control, mediated by a concomitant inhibition of sumoylation and miRNA-directed repression, would thus ensure a rapid, local and effective depletion of RhoGDIα, and its SUMOylated form, to mount *Shigella* invasion (Figure 7).

**Figure 7.**
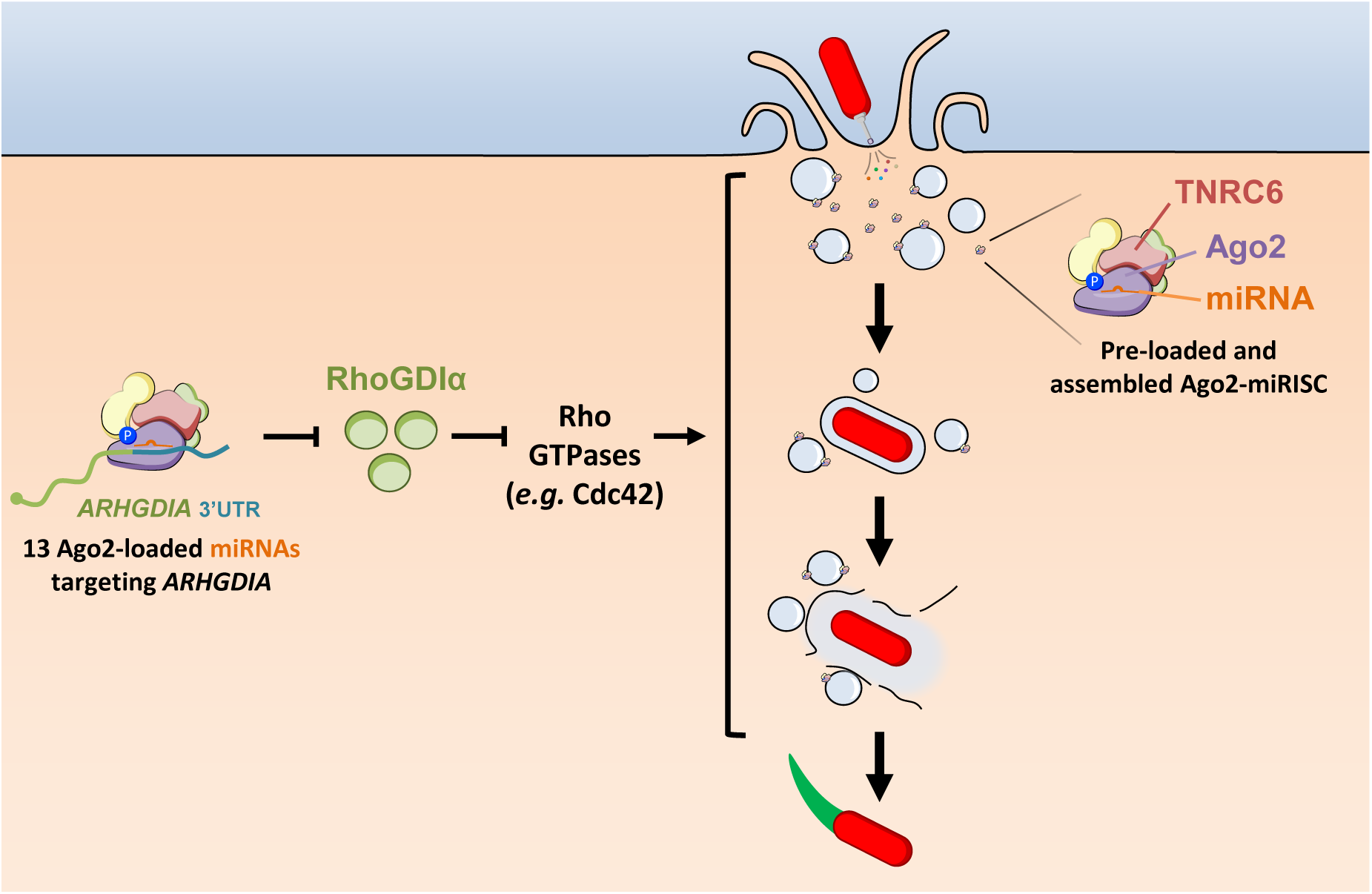
Hypothetical model for the role and regulation of human Ago2 miRISC during early steps of *Shigella* infection. The type III secretion system-dependent internalization of *Shigella* is known to induce membrane ruffling and macropinosome formation^5,10^. *Shigella* is then enclosed in a bacteria-containing vacuole (BCV) whose disintegration is positively regulated by the clustering of infection-associated macropinosomes (IAMs)^10,11^. This facilitates vacuolar escape of *Shigella* to ensure proper bacterial replication and dissemination^13^. In this work, we found that human Ago2 is rapidly and transiently recruited at *Shigella* entry foci. This silencing factor was also found to positively regulate actin foci formation, invasion and vacuolar rupture during *Shigella* infection. None of the expressed miRNAs are differentially regulated or differentially loaded in human Ago2 at 15 min after *Shigella* infection. A mature Ago2-miRISC, characterized by the phosphorylation of Ago2 at serine 387 (blue circle) and the binding of Ago2 partners, like TNRC6 proteins, assembles to form an HMW-miRISC^24,25,30^. This complex is preloaded with miRNAs, including the 13 different miRNAs that target the 3’UTR region of *ARHGDIA* transcripts. Collectively, these miRNAs presumably contribute to the reduced accumulation of RhoGDIα at IAMs^11^ releasing RhoGTPases, like Cdc42^12^, to facilitate vacuolar rupture.

While former studies investigating the role of individual miRNAs in host-microbe interactions have focused on those that are microbe-responsive^19,50^, nothing is known about how miRNAs regulate their targets during immediate steps of infection, prior to their differential expression. Here, we found that none of the human miRNAs are differentially expressed, nor differentially loaded in Ago2, at 15 min after *Shigella* infection. Nevertheless, both the Ago2-dependent miRNA pathway, and the subset of miRNAs targeting *ARHGDIA*, were found to positively regulate vacuolar rupture events. Collectively, these data indicate that the pool of stored miRNAs, pre-loaded in Ago2, is sufficient to regulate this early step of *Shigella* infection. In addition, our work demonstrates that (i) Ago2-S387 phosphorylation, (ii) the ability of Ago2 to bind TNRC6 proteins, and (iii) TNRC6A are all required to promote vacuolar rupture. Because these molecular events, and these Ago2 binding partners, are critical for HMW-miRISC assembly^24–26,30^, our data suggest that an assembled Ago2-mRISC is likely rapidly recruited at *Shigella* entry foci, including IAMs, to orchestrate early steps of infection, and notably vacuolar maturation. Furthermore, because *Shigella* activates the Pi3K-AKT-mTOR signaling pathway^51^, which is known to direct Ago2-S387 phosphorylation^25^, we speculate that its positive effect on vacuolar rupture^11,13^, might be reinforced by promoting Ago2-miRISC assembly. Altogether, these findings suggest that *Shigella* has evolved to hijack the Ago2-dependent miRNA pathway to facilitate vacuolar escape. They also provide evidence that the rapid BCV rupture depends on the miRNA composition of pre-existing Ago2-loaded miRNAs, coupled with Ago2-miRISC assembly. Such regulatory process has previously been reported in mitogen-stimulated quiescent naïve B and T cells, which rapidly re-enter the cell cycle upon activation of Pi3K-AKT-mTOR signalling^52,53^. The above miRNA-dependent mechanism is thus relevant beyond bacterial infection and might be well suited to rapidly silence specific genes in the context of stress responses.

Interestingly, Ago2 predominantly localizes in the nucleus of mouse splenocytes, which are maintained in a quiescent state by the dampening of Pi3K-AKT-mTOR signalling^53^. This results in the co-transcriptional silencing of mobile transposable elements (TEs), including young LINE-1 and ERV/LTR, through Ago2’s catalytic activity^53^. In contrast, stimulating the Pi3K-AKT-mTOR-dependent signalling pathway not only mediates the re-entering of these quiescent cells in the cell cycle, but also restores the cytosolic localization of Ago2, and its miRNA-dependent regulation, through miRISC assembly, thereby releasing the Ago2-dependent silencing of those TEs in the nucleus^53^. It is also noteworthy that gain-of-function mutations in the Pi3K-AKT-mTOR pathway have been associated with colorectal cancers^54,55^, suggesting that a tight control of this signalling pathway is required for intestinal homeostasis. Whether *Shigella*-induced activation of the Pi3K-AKT-mTOR signalling pathway in quiescent cells provides a time window during which young TEs could be derepressed/mobilized, and likewise alter intestinal tissue maintenance and/or orchestrate cancer cell-initiation through their mutagenic effects^56–58^, is an intriguing hypothetical scenario that requires further investigations.

## Material and Methods

### Human cell lines

Human epithelial HeLa cells (ATCC® CCL-2™), CRISPR-Cas9 control (CTL) and knock-out *Ago2*^-/-^ cells^35^ were cultured in DMEM GlutaMAX containing 1.0 g. L^−1^ glucose (Thermofisher) supplemented with 10% fetal bovine serum at 37°C, 5-10% CO_2_. All cell lines were tested negative for mycoplasma contamination across the study.

### Bacterial strains and growth condition

The invasive *Shigella flexneri* 5a wild-type M90T strain^59^ was used in this study. The bacteria were transformed with a pIL22 plasmid expressing the afimbrial adhesin AfaE-1 of uropathogenic *E. coli* to synchronize infection. For microscopy experiments, the M90T strain was transformed with pIL22 derivatives constitutively expressing mCherry or Cerulean (this study). All *Shigella* strains were plated on Tryptic Soy Broth (TSB) agar with 0.01 % Congo red and grown at 37 °C. Bacterial cultures were grown at 37 °C with shaking in TSB (BD, 211825). *E. coli* Top10 strain (Invitrogen) was used for cloning. Bacteria were plated on Lysogeny Broth (LB) Lennox agar and grown at 37 °C. Bacterial cultures were grown at 37 °C with shaking in LB Lennox (BD, 240230). Ampicillin and kanamycin were used at a final concentration of 100 µg. mL^-1^ and 50 µg. mL^-1^, respectively.

### Expression plasmids and constructs

All plasmids and primers used in this study are listed in Tables S2 and S3, respectively. The fluorescent pIL22 derivatives (pMBF5) were generated by cloning 4 fragments (ampR-prom, ampR, *colE1-rop* and *afaI*) amplified from pIL22 and a fluorescent cassette. For the Cerulean, the *cerulean* gene in the same context (Ter-*rpsM*-*cerulean*-Ter) was amplified from pTSAR1Ud2_1^60^, subcloned into a pCR blunt vector (Invitrogen) and cut out using *Not*I/*Spe*I enzymes. For mCherry, the *cerulean* gene from the above-mentioned plasmid was replaced using *Nhe*I/*Hind*III enzymes by the *mcherry* gene from pSU18-mcherry (kind gift of F.-X. Campbell-Valois). The point mutations on *Ago2* were carried out using site-directed mutagenesis by PCR, with primers containing the desired nucleotide changes and the recombinant plasmids selected with *Dpn*I digestion. The *ARHGDIA* gene and 3’UTR were amplified from HeLa cell cDNAs. The *ARHGDIA* was cloned into m3Cerulean-C1 plasmid using *Sac*I and *Bam*HI restriction enzymes. The 3’UTR was further cloned in the *Bam*HI site of the pCFP-*ARHGDIA* plasmid. The pCMV-d2eCFP was generated by replacing the *GFP* gene from pCMV-d2eGFP-miR21^46^ by *CFP* gene from m3Cerulean-C1 plasmid, using specific primers and Gibson assembly. The miRNA sponges that bind the 13 miRNAs targeting *ARHGDIA* transcripts were assembled as described by Lindner *et al.*^61^. Briefly, the 13miR cassette was assembled by concatemeric PCR using overlapping primers. The concatemers were further amplified with three different primer pairs to give cassettes “*Xho*I-D”, “D’E” and “E’-*Apa*I”, with D/D’ and E/E’ corresponding to *Bsa*I sites with complementary 4nt overhangs. The 3 cassettes were further assembled by Goldengate cloning using *Bsa*I. The 3x13miR cassette was finally amplified by PCR and cloned in pCMV-d2eCFP using *Xho*I and *Apa*I restriction enzymes. All constructs were confirmed by Sanger sequencing.

### Bacterial infections

Overnight precultures were diluted to 1/100 into fresh TSB medium and grown at 37 °C with shaking until they reach the exponential growth phase (OD_600_ 0.6-0.8). For live and fixed imaging, bacteria were washed twice with EM buffer (120 mM NaCl, 7 mM KCl, 1.8 mM CaCl_2_, 0.8 mM MgCl_2_, 5 mM glucose, 25 mM HEPES, pH 7.3) before proceeding with cellular infection. For immunoprecipitation experiments, bacteria were washed once in DMEM without serum before proceeding with infection at a multiplicity of infection of 10 for 15 min.

### Transfection conditions

Cells were seeded on glass coverslips into 6-well plate or in 35mm µ-Dish (Ibidi) at a density of 1.5×10^5^ and 2×10^5^ per well 24h prior to targeting siRNA or plasmid transfection, respectively. Cells were transfected the next day with targeting siRNA using Lipofectamine RNAi max (Life Technologies) for 48h or with expression plasmids using JetPrime reagent (Polyplus) for 24h, according to the manufacturer’s instructions. The transfected plasmids and siRNAs are listed in Tables S2 and S4, respectively. For Ago2 complementation experiments, transfection was performed with 0.5 µg GFP, 0.5 µg GFP-Ago2 (or 1 µg where indicated), 1 µg GFP-Ago2^D597A^, ^K660A^, ^Y698A^ and 0.5 µg GFP-Ago2^S387A^, ^S387E^ per well in 6-well plate. All siRNAs were purchased either from Dharmacon (ON-TARGETplus SMARTpool), Thermofisher (Silencer® Select) or QIAGEN (FlexiTube). siRNA knock-down efficiencies were assessed by RT-qPCR or by Western Blot.

### Real time quantitative PCR (RT-qPCR)

Total RNAs were extracted using Quick RNA miniprep kit (Zymo Research) and cDNA prepared with qScript cDNA SuperMix (Quantabio). Real time qPCR was performed using Takyon No ROX SYBR Green master mix (Eurogentec) on a Lightcycler 480 (Roche) instrument. Gene expression was analyzed using 2^(-ΔΔCt)^ methods and normalized to endogenous GAPDH. All primers used are listed in Table S3.

### Western blotting

After siRNA transfection (24h or 48h), cells were washed with PBS and scraped into 1X Laemmli sample buffer. Western blot analysis was performed using the primary and secondary antibodies listed in Table S5. Immunoreactive bands were detected by chemiluminescence using the ECLplus reagent (GE Healthcare Biosciences) and a FujiFilm LAS 4000 system.

### Live cell imaging

Cells were washed twice with EM buffer before proceeding to live cell imaging. To assess vacuolar rupture kinetics (at Collège de France, CdF), living cells were imaged using a wide-field DMIRBE microscope (LEICA microsystems) equipped with a 63x/1.25 oil objective and a Cascade 512 camera (Roper Scientific) driven by the Metamorph (v7.7) software. Every 30 s, a stack of 7z planes (step size of 1 μm) was acquired sequentially for 75 min using a 470 and 546 nm LED source excitation. After acquisition of four time points, bacteria were added at a MOI of 50. At the Institut de Biologie de l’ENS (IBENS), living cells were imaged with a Nikon Ti PFS microscope equipped with a 63x/1.4 oil objective, a CSUX1-A1 (Yokogawa) spinning disk scan head, and an Evolve camera, controlled by the Metamorph (v7.6) software. Every 30 s, a stack of 12 z planes (step size of 0.7 µm) was acquired sequentially in two or three channels for 75 min using a 405, 491 and 561 nm laser source excitation. M90T-mCherry bacteria were added after two time points at a MOI of 30. Maximum projections of each z series were made with ImageJ software, and visually analyzed to determine the time interval between the start of the entry foci formation and the vacuolar rupture.

To study Ago2 recruitment (CdF), transfected cells were also stained for some conditions with the plasma membrane dye CellMask Deep Red Plasma Membrane (DRPM) Stain (Thermofisher) for 10 min in DMEM, 10% FBS at 1:1000 dilution at 37°C, with 10% CO_2_. Cells were imaged with a Zeiss Observer Z1 microscope equipped with a 63x/1.4 oil objective, a CSU-W1 (Yokogawa) spinning disk scan head, and a CMOS (Flash4) camera, controlled by the Metamorph (v7.7) software. Every 30 s, a stack of 22 z planes (step size of 0.4 µm) was acquired sequentially in three channels for 60 min using a 488, 561 and 642 nm laser source excitation. At IBENS, cells were imaged with the aforementioned microscopy system. Every 20 s, a stack of 16 z planes (step size of 0.4 µm) was acquired sequentially in three channels for 60 min using a 491, 561 and 640 nm laser source excitation. For actin recruitment, transfected cells were imaged with the Zeiss system at CdF. Every 30 s, a stack of 21 to 26 z planes (step size of 0.5 µm) was acquired sequentially in two channels for 60 min using a 488 and a 561 nm laser source excitation]. In all settings, after two time points, M90T-mCherry bacteria were added at an MOI of ∼30. Images were analyzed using the ImageJ software. The fluorescence intensities in the emission channels for the proteins of interest were measured within equally sized ROIs around the *Shigella* invasion sites. For each region, the change of slope in the intensity curve in the 488 or 642 (or 640) nm channel was used to determine the time of focus appearance. To determine the fold increase of fluorescent signal at foci, intensity data were normalized by subtracting the background intensity from each time point’s fluorescence intensity (using an area without cells) and then dividing by the fluorescence intensity at T_0_ (corresponding to the average value of the first five time points (about 2.5 min) before foci appearance). Fold increase of normalized fluorescence intensities were represented as curves of the median of the calculated data plotted over time or as median of the maximal fold increase of each foci.

### Fixed cell imaging

Cells were seeded on glass coverslips into 24-well plate the day before infection. The next day, cells were washed twice with EM buffer, incubated with the DRPM stain (Invitrogen) and washed three times before proceeding to infection. M90T-mCherry bacteria were added at a MOI of 30 and incubated for 30 min at 37°C + 5% CO_2_. Infected cells were washed three times with PBS, fixed for 20 min with PFA 4%, washed three times and finally, PFA was quenched with 0.1M glycine for 10 min. After three washes, extracellular bacteria were labelled with mouse anti-LPS^62^ antibodies in PBS + 10% FBS for 1h at RT. Coverslips were then washed three times and incubated for 1h with anti-mouse Alexa488 in PBS +10% FBS. Coverslips were finally washed three times and mounted on ProLong™ Gold Antifade Mountant (Invitrogen). Images were acquired with a Zeiss Observer Z1 microscope equipped with a 63x/1.4 oil objective, a CSU-W1 (Yokogawa) spinning disk scan head, and a CMOS (Flash4) camera.

### RNA-immunoprecipitation

Immunoprecipitation of Ago2 was performed on semi-confluent HeLa cells cultured in 15-cm plates as described by Rüdel *et al.*^63^. Briefly, 16 µg of anti-Ago2 antibodies were bound to Dynabeads Protein G (Thermofisher) at 4 °C. Control and infected cells were washed with PBS and scrapped in lysis buffer (Tris-HCl 20 mM, pH 7.4; NaCl 150 mM; EDTA 2 mM; NP-40 0.5 %; DTT 0.5 mM) containing complete protease inhibitor (Sigma-Aldrich) and RNAseOUT (Thermofisher). One tenth of the cleared lysates were mixed with 500 µL TRIzol (Thermofisher) and kept at -80 °C for further RNA extraction. The lysates were incubated overnight at 4 °C with the anti-Ago2 coated beads. After three washes with IP wash buffer (Tris-HCl 50 mM, pH 7.4; NaCl 300 mM; NP-40 0.05 %) containing RNaseOUT, one tenth of the beads were mixed with Laemmli buffer for western blotting and the remaining beads were incubated with 20 µg Proteinase K. The eluates were mixed with 1 mL TRIzol and both input and eluate RNAs were extracted according to the manufacturer’s instructions (Thermofisher). The RNA extracts were processed and small RNAs sequenced by Fasteris (Switzerland).

### Ago2-bound miRNAs identification through small RNA data mining

Small RNA reads were mapped against the human genome (GRCh38.p7), obtained from NCBI, using Bowtie2 (v2.3.2, --end-to-end mode)^64^. Annotated miRNA coordinates were obtained from miRBase (v21)^65^ and counts of mapped reads for annotated miRNAs were obtained using HTSeq-count^66^. Normalized counts and differential accumulation of miRNAs in libraries was assed using DESeq2^67^. Data on experimentally validated microRNA-target pairs were obtained from miRTarBase^68^ and miRWalk^69^. Gene Ontology term enrichment was assessed using clusterProfiler in R^70^.

### Statistical analyses

All statistical tests were performed using the GraphPad software (v6). Given the non-Gaussian distribution of foci formation, Ago2 recruitment and vacuolar rupture timing, differences were assessed using the Wilcoxon Mann Whitney test for the comparison of two groups and the Kruskal-Wallis one-way ANOVA when more groups were compared. The Chi-square (χ^2^) test was performed to compare the frequency distribution of early (≤ 4 min) *vs.* later (> 4 min) ruptures.

## Acknowledgments

We thank Pr. Gunter Meister for providing the YFP-T6B and -T6B mutant peptides, and Dr. François-Xavier Campbell-Valois for providing the pSU18-mcherry plasmid. We kindly thank Dr. Yuen-Yan Chang for her help with the experiments presented Figures S3C and S3D, as well as Dr. Jost Enninga, for its valuable criticisms on the first version of this manuscript. We acknowledge members of the Navarro laboratory for critical reading of the manuscript. Finally, we thank the IBENS imaging facility for providing access to the microscopy equipment that played a key role in this work and for providing assistance whenever necessary. This work was supported by a grant from the European Molecular Biology Organization (EMBO) Young Investigator Program (to L.N.), an Emergence grant from the Mairie de Paris (to L.N.), a grant from DIM Malinf (to L.N.), a Grant from the Bettencourt Schueller Foundation (to L.N.), a grant from the Agence Nationale de la Recherche (ANR-10-LABEX-54 MEMOLIFE) (to L.N. and G.T.V.N.).

## Author contributions

DF, LS, MB, LN, designed research; FD, LS, MB, MC, performed research; A P-Q, conducted bioinformatic analyses; FD, LS, MB, A P-Q, BS, PS, LN, analyzed data; DF, LS, MB, LN, wrote the manuscript; G.T.V.N, LN, acquired fundings; all the authors reviewed and edited the manuscript.

## Competing Interest Statement

All the authors declare no competing interests.

